# ProteoScopeR: A Shiny Workflow for Method Comparison and Reproducible Quantitative Proteomics Analysis

**DOI:** 10.64898/2026.09.20.752977

**Authors:** Benbo Gao, Hanqing Zhao

**Affiliations:** Biomedical Research Center, Sir Run Run Shaw Hospital, Zhejiang University School of Medicine, Hangzhou, Zhejiang 310016, China; Beijing Tongren Eye Center, Beijing Tongren Hospital, Capital Medical University, Beijing, China

**Keywords:** quantitative proteomics, method comparison, normalization, missing-value imputation, differential abundance, Shiny, reproducibility

## Abstract

Quantitative proteomics requires decisions about normalization, missing-data handling, and statistical modeling that can change the reported results. ProteoScopeR is an R package and Shiny application that connects these decisions in a traceable workflow and complements downstream exploration in xOmicsShiny. Side-by-side comparisons expose changes in distributions, feature retention, effect estimates, and selected protein sets. An optional external artificial-intelligence assistant reviews exported evidence and proposes settings for researcher approval; statistical calculations remain in R. A group-only case study used DIA-NN-derived aqueous humor data from 69 samples and 3,667 imported protein groups. Six normalization methods and seven missing-data strategies were screened, and the approved cyclic-loess analysis compared five differential workflows. For nAMD versus pmCNV, limma and proDA selected 54 and 37 protein groups, respectively, with 31 shared selections at adjusted P < 0.05 and absolute log_2_ fold change ≥ 0.5. Input routing, scientist decisions, locked settings, and an execution receipt preserve the analysis record. These comparisons describe sensitivity to analytical choices rather than identify a universally superior method.

## Introduction

Quantitative proteomics is widely used to study molecular changes in disease, experimental perturbations, and biological development. Identification and quantification produce an abundance table, but several analytical decisions remain before that table can support a biological conclusion. Sample annotations need to match the measurements, statistical comparisons should reflect the experimental design, and preprocessing has to accommodate variation and missing values in the data. These steps can be difficult to inspect when they are spread across separate scripts and applications.

We previously developed Quickomics to support interactive exploration of analyzed omics data.^1^ Its successor, xOmicsShiny, extends this approach to cross-omics comparisons, pathway mapping, networks, and expression trends.^2^ Both benefit from carefully prepared abundance matrices, sample annotations, and statistical results. ProteoScopeR is a companion R package that supplies the proteomics preprocessing and statistical analyses preceding that exploration.

Several applications address parts of this workflow. MatrixQCvis provides interactive quality assessment of matrix-like omics data, DEP2 combines proteomics analysis with biological interpretation, and MSstatsShiny provides a graphical interface to the MSstats family.^3-5^ ProteoScopeR connects design checks, preprocessing comparisons, differential analysis, and optional AI-assisted review in one interface. Its exports retain the settings and matrices used in each analysis, making the path to the results easier to inspect. Table 1 places this scope alongside related applications.

**Table 1.** Scope of ProteoScopeR and related applications.

| Application | Main analytical scope | Typical output |
| --- | --- | --- |
| Quickomics<sup>1</sup> | Interactive exploration of analyzed omics data | Figures and result tables |
| xOmicsShiny<sup>2</sup> | Cross-omics comparison and biological interpretation | Integrated views, pathways, and networks |
| MatrixQCvis<sup>3</sup> | Quality assessment and preprocessing of omics matrices | QC reports and processed objects |
| DEP2<sup>4</sup> | Quantitative proteomics analysis and post-analysis | Statistical and biological results |
| MSstatsShiny<sup>5</sup> | MSstats-family processing and statistical analysis | MSstats results and reports |
| ProteoScopeR | Design checks, preprocessing comparisons, multiple differential-analysis engines, and optional recorded AI review | Comparison figures, statistical tables, reusable objects, and recorded decisions |

The table summarizes the principal scope of each application; it is not an exhaustive feature inventory or a performance comparison.

Here, we describe ProteoScopeR and illustrate its use with experimental aqueous humor proteomics data. Method comparison is a central function of the application: users can inspect how alternative normalization, missing-data, and differential-analysis choices affect the same dataset. Linking these comparisons to their input matrices, model specifications, and thresholds supports transparent method selection and helps distinguish stable findings from results that depend on a particular workflow. The application starts from quantified features and produces statistical tables and reusable R objects, with optional AI-assisted evidence review and a dedicated export for xOmicsShiny.

## Experimental Section

### Software Architecture and Data Import

ProteoScopeR is an R package with eight Shiny pages (Figure 1) and standalone functions for import, design construction, preprocessing, statistical fitting, and export. Its native data object stores the assay, annotations, source information, processing history, design, matrices, settings, and results. SummarizedExperiment and QFeatures conversions support exchange with Bioconductor workflows.^6^ Optional dependencies are checked when a method is requested; unavailable methods and fitting failures are reported rather than replaced by a different statistical engine. Supporting Information Table S1 summarizes inputs, operations, and outputs.

**Figure 1.**
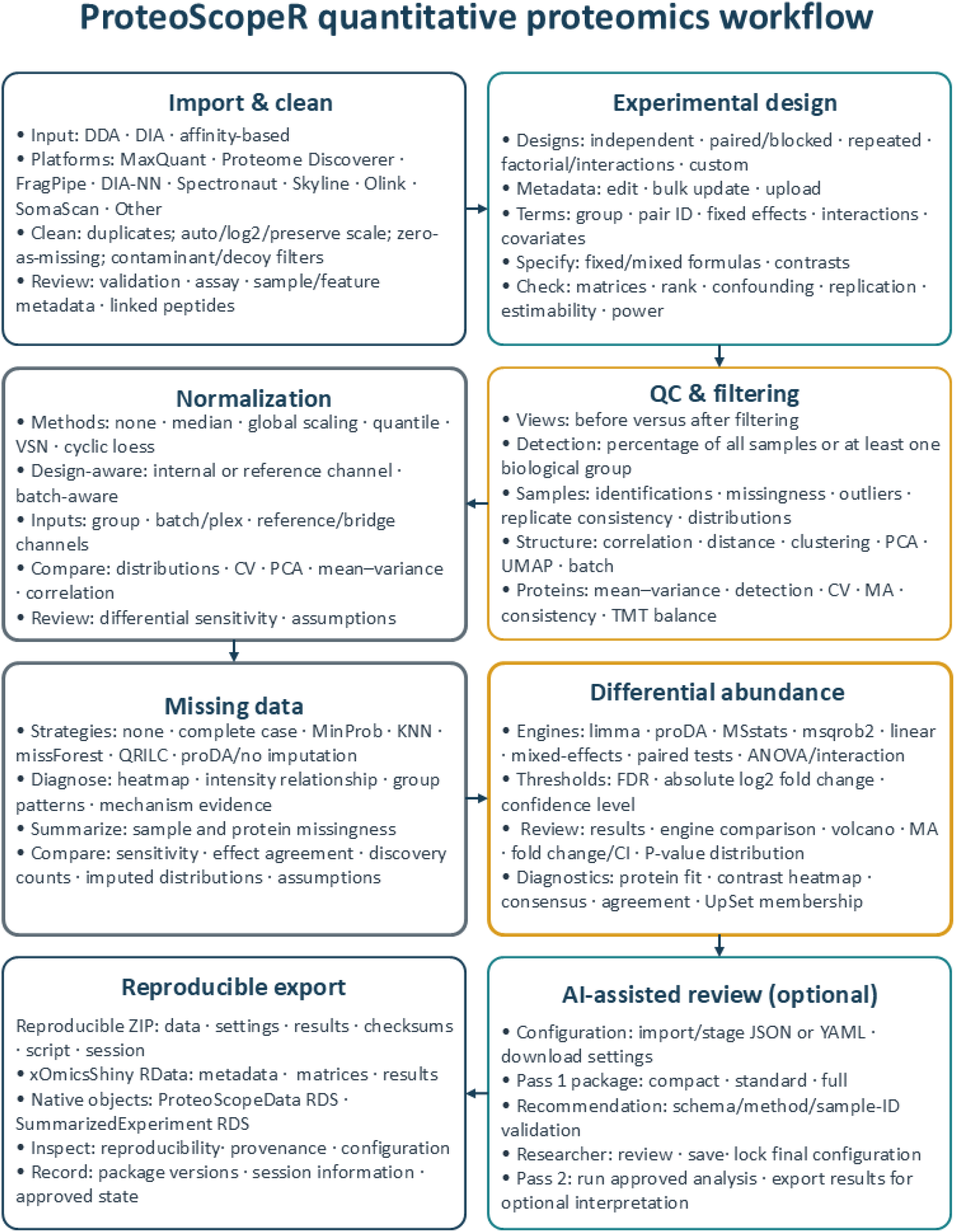
ProteoScopeR quantitative proteomics workflow. Eight connected pages cover import and validation, experimental design, quality control and filtering, normalization, missing-data evaluation, differential analysis, optional AI-assisted review, and reproducible export. Method comparisons are embedded in the normalization, missing-data, and differential-analysis stages; their settings and outputs support researcher review before final analysis choices are recorded.

The input page lists the files required for the selected acquisition method and processing platform; users select the acquisition method and processor explicitly, while file-format and column-schema recognition remains available for generic programmatic imports. For DIA-NN, users provide protein-group and precursor matrices plus optional sample metadata. Protein groups form the analysis assay, while precursor measurements are retained for traceability and quality assessment. Other routes support MaxQuant, Proteome Discoverer, FragPipe, Spectronaut, Skyline, Olink, SomaScan, manually mapped tables, and supported R objects. Checks cover identifiers, quantitative columns, contaminants and decoys, duplicate entries, value scale, nonfinite values, sample names, and metadata alignment.

### Experimental Design and Quality Assessment

Users define grouping variables, model formulas, and contrasts from the sample metadata. Supported designs include independent, paired or blocked, repeated-measures, and factorial models. ProteoScopeR checks model rank, residual degrees of freedom, confounding, replication, and contrast estimability; these checks do not determine whether every biologically relevant covariate has been included.

Quality assessment covers identification counts, missingness, abundance distributions, correlations, principal component analysis, clustering, coefficients of variation, and outlier diagnostics. Batch associations and assay-specific checks are available when the required information is present. Detection filters can be applied across all samples or within groups, and the rule and retention counts are recorded.

### Normalization, Missing Data, and Differential Abundance

Normalization options include retaining imported values, median normalization, global scaling, and quantile normalization that accommodates missing entries. Variance-stabilizing normalization, cyclic loess, reference-based procedures, and batch-aware methods are available where their requirements are met.

Candidate methods are compared using shared-axis abundance distributions, within-group variation, sample correlations, and available batch diagnostics. The numerical screen includes a within-group coefficient of variation calculated after exponentiating analysis values; this metric is scale-dependent, particularly after VSN, and is not an accuracy criterion. Distribution alignment and lower variation must be considered alongside biological context.

Missing-data options include no imputation, complete-case selection, MinProb, K-nearest neighbors (KNN), missForest, and QRILC. MinProb delegates to imputeLCMD::impute.MinProb with default q = 0.01 and tune.sigma = 1.^14^ Missingness patterns cannot uniquely establish the generating mechanism.^8^ ProteoScopeR therefore compares retained features, remaining missingness, filled-value distributions, and changes in screening effects. The no-imputation/proDA option specifies a downstream dropout-aware model, not a separate imputation algorithm.^9^ The pre-imputation matrix is retained for engines configured to receive incomplete observations.

Normalization and missing-data screens use the first two biological groups in metadata order to calculate mean differences and feature-wise Welch tests, followed by BH adjustment, using adjusted P < 0.05 and an absolute log_2_ fold-change cutoff of 0.5. These exploratory screens do not use the full configured model. Even in a group-only analysis, a two-group Welch screen and a model fitted across all four groups need not produce the same selections.

Differential-abundance options include limma, proDA, MSstats, msqrob2, mixed-effects models, and base R analyses appropriate to the design.^7,9,10,13^ Results share feature and contrast identifiers, method labels, effects, uncertainty, P values, adjusted P values, and issue flags. The comparison interface displays shared-axis volcano plots, effect agreement, and selected-set intersections. Users may restrict plots to proteins with finite effects and adjusted P values in every engine, while preserving each engine’s original multiple-testing adjustment. Issue flags distinguish non-estimable tests from unselected proteins. The input matrix, imputation, covariates, and testable universe must be matched before differences can be attributed to model choice alone.

### Optional AI-Assisted Review

AI review uses manual file exchange; no integrated model API or automatic external transmission is required. Pass 1 exports instructions, a response schema, an example, design diagnostics, QC, and available method comparisons. The researcher may send this package to an external service and import its JSON or YAML recommendation. ProteoScopeR validates field types, run-specific method choices, duplicate sample flags, and the review identifier. Scientist rationale is retained and required for changed recommendations and flagged-sample decisions; flagged samples default to retention. The researcher locks the selected settings and runs the approved analysis, which rebuilds the design and reruns filtering, preprocessing, and statistical fitting. Pass 2 exports those results for optional interpretation. Both AI exports offer pseudonymization and metadata minimization, not anonymization; users must inspect the contents before external transfer. Model provenance and examples are described in Supporting Information.

### Reproducible Outputs and xOmicsShiny Export

General outputs include statistical tables, annotations, native R objects, and a reproducibility bundle containing parameters and intermediate matrices. Bundles preserve validation summaries, available comparisons, checksums, package and session information, and a reproduction script. Approved runs additionally retain the locked configuration, decision audit, and execution record. Final downloads use the saved approved state and reject stale results. The ProteoScopeR replay script checks package identity, engine completeness, result keys, and numerical agreement; it starts from the saved imported object rather than raw mass spectra.

The dedicated xOmicsShiny RData export contains six objects: MetaData, ProteinGeneName, data_wide, data_long, results_long, and data_results. They supply sample annotations, feature mappings, abundance values, per-test statistics, and combined result summaries. Feature and sample identifiers are checked for alignment before export, and test names include the statistical method to distinguish parallel analyses. In data_wide, nonfinite entries are replaced with zero to meet the downstream format; the analysis matrices preserve their own missing-value representation. This conversion is an export convention and should not be interpreted as an abundance estimate.

### Experimental Data and Case-Study Analysis

The case study used aqueous humor measurements from the previously described clinical cohort.¹¹ Sample collection, liquid chromatography–mass spectrometry, and DIA-NN identification and quantification are described in the source study; its biological conclusions were not used as a reference standard.¹¹,¹² The imported dataset contained 3,667 protein groups from 69 samples: 21 with neovascular age-related macular degeneration (nAMD), 20 with pathologic myopia–related choroidal neovascularization (pmCNV), 20 age-related cataract controls (Ctrl), and eight with pathologic myopia without choroidal neovascularization (PM). The analysis was performed at protein-group level.

The case study used an independent-sample design with formula ∼ 0 + group and three contrasts: nAMD– pmCNV, pmCNV–PM, and nAMD–Ctrl. Age and sex were excluded from every configured model so the same group-only specification could be used with the MSstats. Detection in at least 30% of all samples retained 3,662 protein groups and removed five; all 69 samples were retained.

Six normalization candidates were screened: none, median, global scaling, quantile, VSN, and cyclic loess. The seven missing-data strategies were compared after cyclic loess normalization, matching the normalization selected for the final run. The case-study exports retain the imputation choices used to generate each comparison, which are stated below. These comparisons are sensitivity analyses tied to the specified method and do not establish a universally preferable imputation strategy.

The five differential workflows comprised limma, proDA, MSstats, msqrob2, and an unmoderated linear model. limma and the linear model received the complete canonical-MinProb-imputed matrix generated by imputeLCMD::impute.MinProb (q = 0.01, tune.sigma = 1). proDA, MSstats, and msqrob2 received the same cyclic-loess-normalized matrix with 43,836 missing entries. All used the group-only specification; the MSstats represents each protein group as a synthetic feature and is not a precursor-level MSstats workflow. Table S2 records matrix routing, model specification, and testing universes; Table 2 and Supporting Information Section S3 report testable and selected counts. The preprocessing screens and final fits remain separate computational stages.

**Table 2.** Selected and testable protein groups in the group-only analysis at adjusted P < 0.05 and |log_2_ fold change| ≥ 0.5.

| Engine | nAMD–pmCNV | pmCNV–PM | nAMD–Ctrl |
| --- | --- | --- | --- |
| limma | 54 / 3,662 | 3 / 3,662 | 20 / 3,662 |
| proDA | 37 / 3,662 | 0 / 3,662 | 4 / 3,662 |
| msqrob2 | 113 / 3,662 | 10 / 3,659 | 57 / 3,661 |
| MSstats | 68 / 3,662 | 4 / 3,659 | 23 / 3,661 |
| Linear model | 54 / 3,662 | 2 / 3,662 | 25 / 3,662 |

The case study used the default selection criteria: finite log_2_ fold changes and adjusted P values, adjusted P < 0.05, and absolute log_2_ fold change ≥ 0.5. The confidence level for interval estimates was 95%. Adjusted P values were preserved within each engine–contrast analysis and were not recomputed after selecting common-testable proteins. An independent ordinary least-squares calculation in the accompanying evidence package checked the exported linear-model coefficients, standard errors, and P values against the saved matrix and design.

## Results and Discussion

### Quality Assessment and Feature Retention

The group-only design had full rank, 65 residual degrees of freedom, and estimable contrasts. Group sizes ranged from 8 to 21. The imported assay contained 44,086 missing entries among 253,023 possible measurements (17.4%). Following detection filtering, 43,836 of 252,678 entries were missing (17.35%); sample-level missingness ranged from 10.87% to 30.64% (Figure 2A). Complete-case selection retained 1,025 of 3,662 filtered groups (27.99%) and discarded 2,637 (Figure 2B). These are successive feature universes, not conflicting counts.

**Figure 2.**
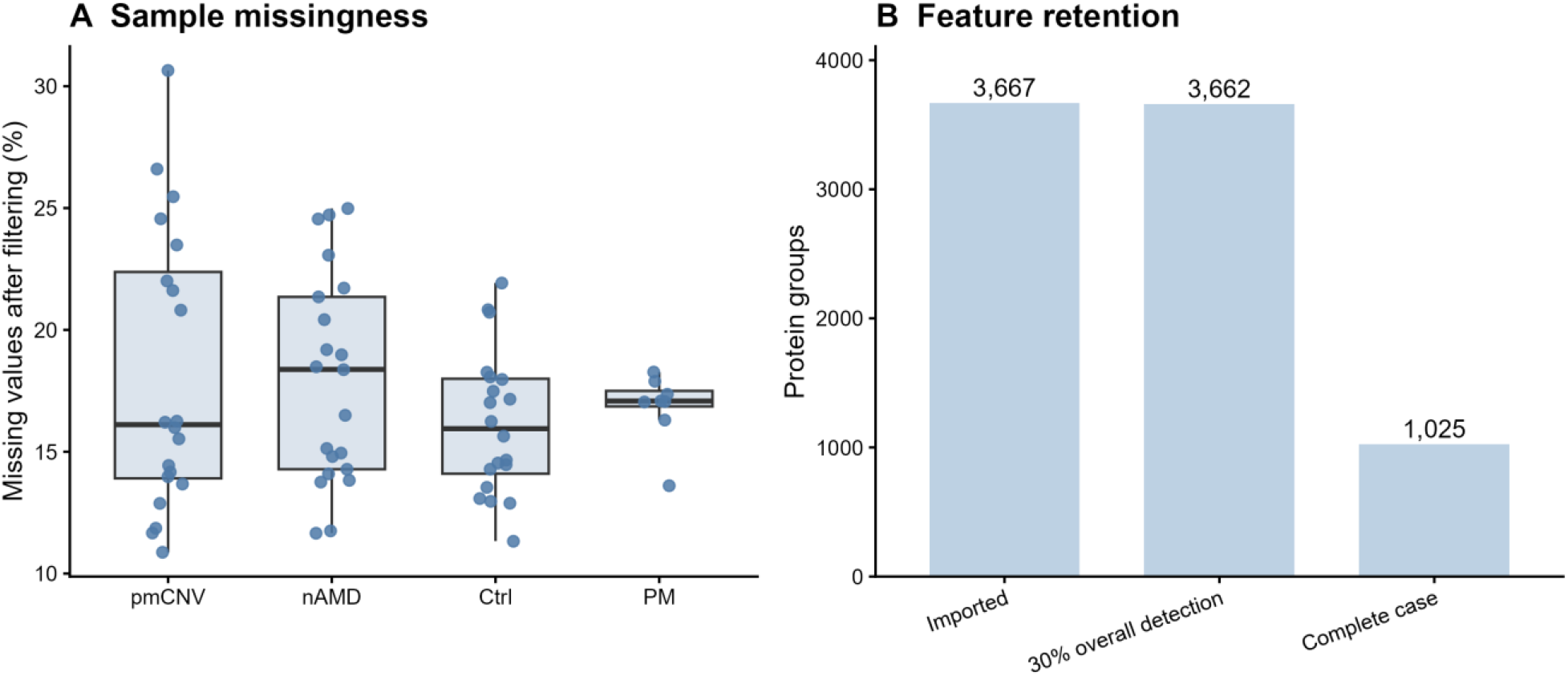
Quality assessment and feature retention in the group-only case study. (A) Sample missingness after 30% overall-detection filtering. Points are samples; boxes show medians and interquartile ranges, with whiskers extending to values within 1.5 interquartile ranges. (B) Protein groups at import, after detection filtering, and under complete-case selection. Values were generated from the analysis matrices.

Input QC flagged four samples for review, and the imported AI recommendation proposed investigation rather than automatic exclusion. The approved configuration retained all four. The decision audit contains the recommendations and final actions.

### Comparison of Normalization Methods

The normalization screen showed small differences in within-group variability and sample correlation (Figure 3). The median within-group CV was 0.4943 without normalization, 0.4943 after median normalization, 0.4951 after global scaling, 0.4918 after quantile normalization, 0.4919 after VSN, and 0.4859 after cyclic loess. Median sample correlations ranged from 0.8825 to 0.8835. Figures S1 and S2 provide additional normalization-screen metrics from the supplied comparison table.

**Figure 3.**
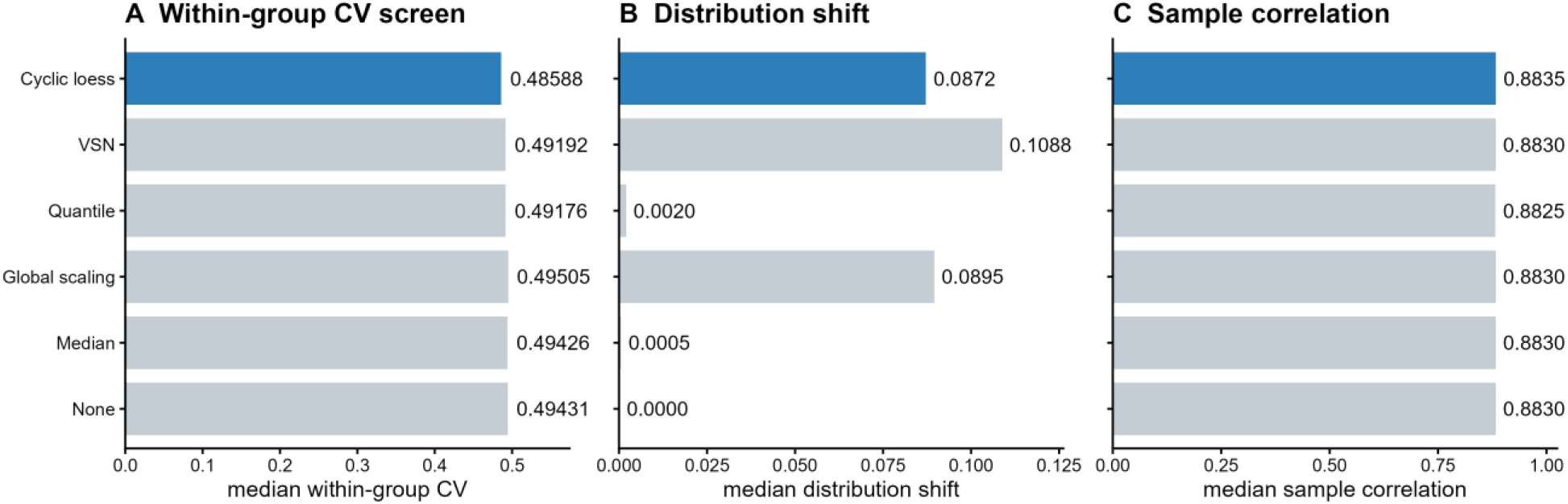
Screening summaries for six normalization methods applied to the same 3,662 filtered protein groups and 69 samples. Panels show (A) median within-group CV, (B) median distribution shift, and (C) median sample correlation. Cyclic loess, selected for the approved run, is blue. *CV is computed after exponentiation of analysis values and is scale-dependent, especially after VSN. These summaries do not rank normalization accuracy.

The median absolute shift in sample medians was 0.0005 for median normalization, 0.0895 for global scaling, 0.0020 for quantile normalization, 0.1088 for VSN, and 0.0872 for cyclic loess. The recommendation favored cyclic loess while acknowledging its greater distributional change relative to median and quantile normalization. This records the decision and its trade-off, rather than a pre-specified or independently validated optimum.

Quick-screen selections ranged from five to 19 across normalization candidates. These counts were not used as an objective to maximize. The comparison instead makes transformation-dependent changes visible before fitting the selected downstream workflows.

### Comparison of Missing-Data Strategies

The seven-strategy screen used cyclic-loess-normalized input (Figure 4). No imputation and no imputation with downstream proDA retained all 3,662 groups and left 17.35% of entries missing. Complete-case selection retained 1,025 groups. MinProb, KNN, missForest, and QRILC retained all 3,662 groups and filled all 43,836 missing entries. Table S3 and Figure S3 summarize the retained features, effect agreement, and filled-value locations reported in the comparison table.

**Figure 4.**
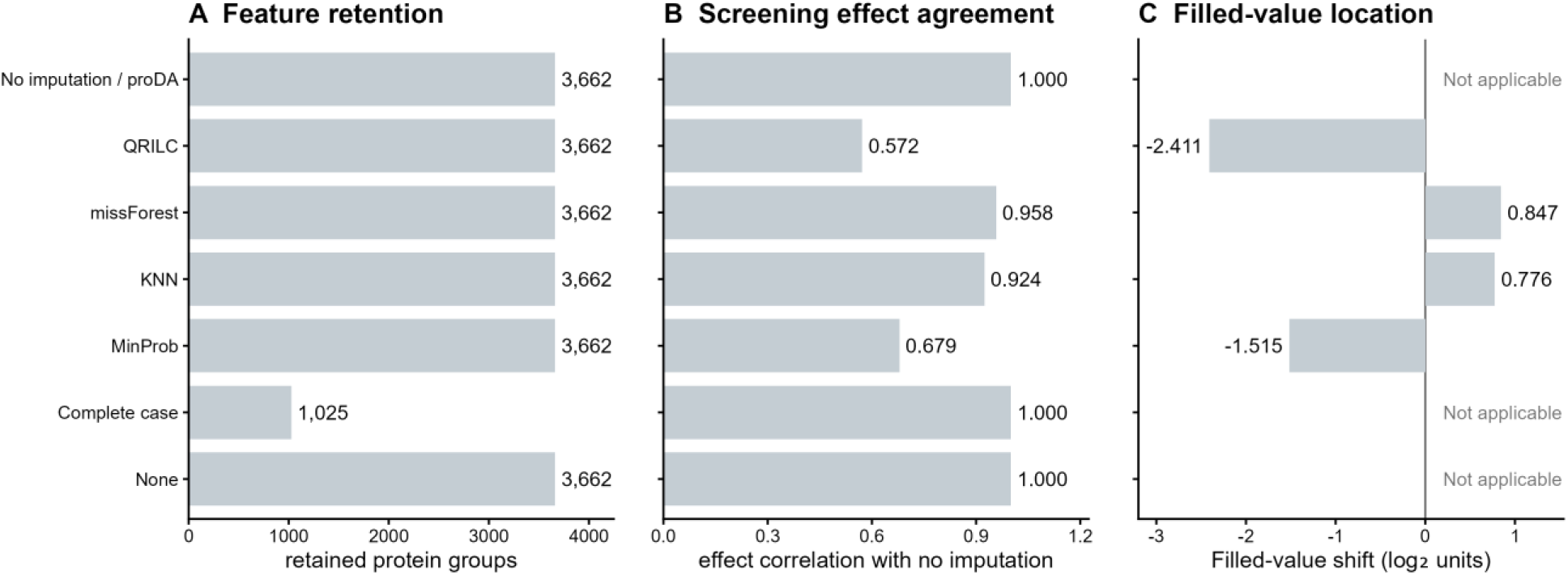
Missing-data sensitivity screen after cyclic loess normalization. (A) Retained protein groups. (B) Pearson correlation of nAMD-minus-pmCNV screening mean differences with no imputation, on shared proteins. (C) Median filled value relative to the observed 10th percentile. MinProb denotes imputeLCMD::impute.MinProb with the parameters specified in Methods. No imputation/proDA denotes unchanged input, not a fitted proDA model. Complete-case agreement covers its reduced universe; filled-value locations are not applicable without imputation.

Screening effect correlations with no imputation were 0.679 for MinProb, 0.924 for KNN, 0.958 for missForest, and 0.572 for QRILC. Median filled values lay 1.515 and 2.411 log_2_ units below the observed 10th percentile for MinProb and QRILC, respectively, but 0.776 and 0.847 units above it for KNN and missForest. These locations expose the contrasting assumptions used to reconstruct unobserved measurements.

The no-imputation/proDA entry had a correlation of one because the preprocessing screen did not fit proDA or change the matrix. Complete-case agreement likewise concerned only retained proteins. Quick-screen selections were 13 without imputation, 58 after complete-case selection, 14 after MinProb imputation, 17 after KNN, 15 after missForest, and 13 after QRILC. Complete-case selection changes both the feature universe and the multiple-testing family. Neither a larger selected set nor stronger effect agreement verifies the missing values.

### Differential Analysis and Agreement Across Workflows

The ProteoScopeR analysis bundle contained 54,930 differential-result rows: 3,662 protein groups × three contrasts × five engines. For nAMD–pmCNV, limma selected 54 groups, proDA 37, msqrob2 113, the MSstats 68, and the linear model 54 (Table 2). For pmCNV–PM the corresponding counts were three, zero, ten, four, and two; for nAMD–Ctrl they were 20, four, 57, 23, and 25. The corresponding volcano plots are shown in Figure S4. These are finite-effect selections using both thresholds, not adjusted-P-only counts or pooled unique proteins.

Entries are selected / testable groups. Testability requires finite effect and adjusted P value. All workflows used cyclic loess and the group-only specification. limma and the linear model used the MinProb-imputed matrix; the other three engines used the common normalized incomplete matrix. Original engine–contrast adjustments are retained. Matrix routing and the protein-level MSstats are detailed in Table S2.

The MSstats returned four nonfinite feature–contrast effects: three for pmCNV–PM and one for nAMD–Ctrl, all flagged oneConditionMissing. Their exported adjusted P values were zero despite unavailable raw P values, so these rows were excluded by the finite-effect rule. msqrob2 likewise had three non-estimable tests for pmCNV–PM and one for nAMD–Ctrl. No engine lost testability for nAMD–pmCNV. Diagnostic rows remain in the analysis outputs rather than being converted to zero effects.

All 3,662 groups were testable in all five nAMD–pmCNV workflows (Figure 5). limma and proDA shared 31 selections, with 23 limma-only and six proDA-only; their effect correlation was 0.805. All 37 proDA selections were also selected by both msqrob2 and the MSstats. Correlations were 0.935 for proDA– msqrob2, 0.955 for proDA–MSstats, and 0.979 for msqrob2–MSstats. Thirty-one groups were selected by all five workflows. Full pairwise results are provided in Table S4.

**Figure 5.**
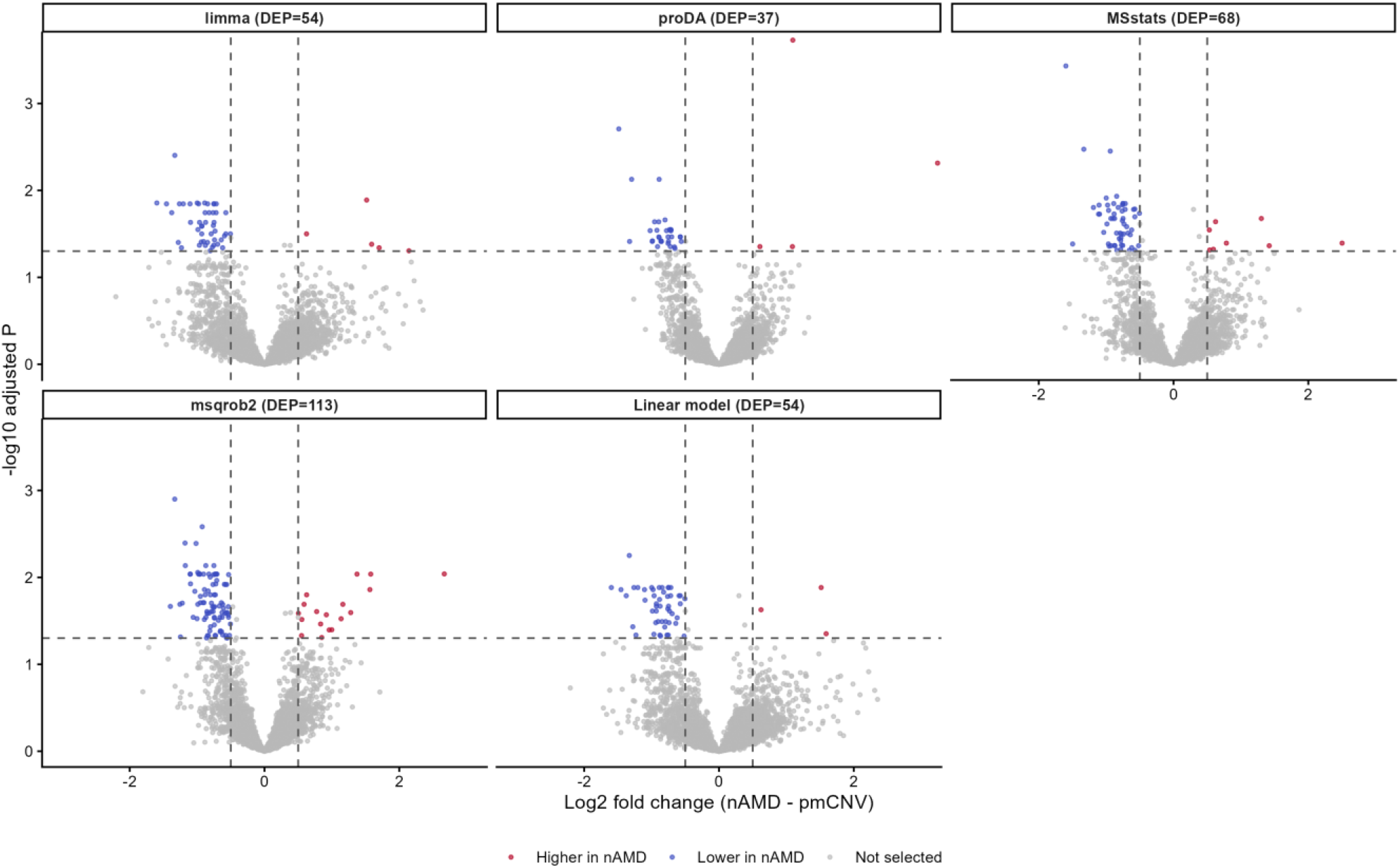
Approved group-only differential workflows for nAMD–pmCNV. All panels use the 3,662 groups testable in every engine and shared axes. Dashed lines mark adjusted P = 0.05 and log_2_ fold change = ± 0.5. Red and blue points pass both cutoffs and indicate higher and lower abundance in nAMD, respectively; gray points do not. limma and the linear model use MinProb-imputed input; proDA, the MSstats, and msqrob2 use the normalized incomplete matrix. Adjusted P values retain their original engine–contrast testing families.

The comparisons include two matched-input subsets. limma and the linear model used identical complete input and had identical effect estimates to numerical precision; each selected 54 groups, with 52 shared selections. proDA, msqrob2, and MSstats used the same incomplete matrix and group-only specification, although their modeling assumptions and the MSstats representation differ. Matching the group-only specification controls covariate specification across workflows; it does not make the imputed and non-imputed workflows identical.

DEP denotes the number of protein groups meeting both selection thresholds in each panel.

### Method Comparison as a Primary Software Output

The case study illustrates complementary comparison tasks: normalization changes distribution summaries, missing-data strategies change retention and reconstructed values, and differential workflows change testability and selected sets. ProteoScopeR presents these comparisons alongside the input population, matrix treatment, model, thresholds, and unavailable results so that the reason for a selection can be recorded.

Independent verification checked the locked version and configuration checksum, input fingerprint, result dimensions, matrix alignment, finite-testable counts, and pairwise intersections. Ordinary least squares on the saved downstream matrix reproduced all 10,986 linear-model tests; the maximum absolute discrepancy in adjusted P values was below 3 × 10−^13^. This confirms the exported linear-model calculations and counting logic, not empirical FDR control, imputation accuracy, or validation of every adapter.

### AI-Assisted Method Review and Interpretation

The imported recommendation favored cyclic loess and no imputation with proDA, while the scientist retained the MinProb-imputed branch and all five engines for comparison. The JSON recommendation, exported instructions, schema, approved settings, and decision audit preserve this difference between advice and execution. The case-study recommendation does not identify its model provider or generation settings; these are reported as not recorded, without attributing them to a model from the manuscript-writing acknowledgments.

The approved analysis package contains the locked configuration, scientist decision audit, execution receipt, QC evidence, results, and cross-engine agreement needed for evidence-linked interpretation. ProteoScopeR records supplied model provenance and asks for no more than three unresolved questions that could materially change a decision. Its schema and validation constrain the response format and allowed actions, but do not verify the truth of narrative evidence or biological claims.

AI review remains optional. Statistical calculations are performed in R, and the scientist is responsible for settings and interpretation. This example documents a review process; it does not measure whether AI assistance improves analytical accuracy.

## Limitations

This single-platform clinical case study demonstrates researcher-guided workflow comparison and provides no ground truth for evaluating imputation or differential-abundance accuracy. Unequal group sizes and age distributions limit interpretation of the deliberately unadjusted comparisons, and the protein-group input used for MSstats does not represent its full precursor-level workflow.

Controlled benchmarks were outside the scope of this workflow demonstration. Such benchmarks provide valuable performance estimates under defined conditions, but findings may not generalize across experimental designs, acquisition platforms, and heterogeneous biological samples. No accuracy, scalability, or AI model-quality benchmark is claimed.

ProteoScopeR presents alternative methods side by side without treating descriptive diagnostics as accuracy rankings or imposing an automated pipeline choice. Researchers select methods according to their experimental design and scientific question. Comparative visualizations, execution receipts, and audit trails support documenting and evaluating how these choices affect results.

ProteoScopeR begins with quantified abundance tables and does not search or reprocess raw mass spectra. Its results depend on upstream identification and quantification, data quality, experimental design, and the assumptions of the selected statistical methods. Method comparisons make these dependencies visible but do not replace validation, domain expertise, or scientific judgment.

## Conclusions

ProteoScopeR complements xOmicsShiny with design checks, preprocessing comparisons, and multi-engine proteomics analysis. The group-only aqueous humor example shows how workflows can be compared under a common group-only specification while differences in matrix treatment, model assumptions, and testability remain explicit. Optional AI review organizes exported evidence into proposals that researchers approve before execution. Retaining the comparisons, original results, settings, and execution record supports inspection and reuse without treating agreement or discovery counts as measures of accuracy.

## Supporting Information

Workflow inputs and outputs; exact case-study configuration and matrix routing; missing-data metrics; testability, selections, and pairwise agreement; AI evidence provenance and safeguards; additional comparison plots. The accompanying evidence package contains the source tables and verification scripts.

## Data and Software Availability

ProteoScopeR is freely available under the MIT License. Source code and documentation are available at https://github.com/bbgao/ProteoScopeR. The accompanying evidence package used for the case study, ProteoScopeR_analysis_bundle_v1.0.0.zip, contains the locked ProteoScopeR v1.0.0 configuration, processed matrices, result tables, and verification scripts. This package is available in the repository under the case-study/results/ directory. The case-study/ directory also contains the corresponding configuration and replay materials. The input files used for the case study, including pg_matrix.tsv, pr_matrix.tsv, and metafile.csv, are provided under case-study/input/. An automated GitHub Actions workflow (R-CMD-check) builds and checks the package on Windows, macOS, and Ubuntu release/devel environments on every push. The source cohort and acquisition methods are described by Zhao et al.^11^

## Author Information

### Author Contributions

Benbo Gao designed the software, developed and maintained the code, and wrote the manuscript. Hanqing Zhao tested the software, contributed to manuscript preparation, and wrote the user manual. B.G. and H.Z. contributed equally to this work.

### Notes

The authors declare no competing financial interest.

## Acknowledgments

During manuscript preparation, the authors used OpenAI ChatGPT and xAI Grok to assist with coding, debugging, software packaging, language editing, and document formatting. The authors reviewed and revised all generated content and take full responsibility for the manuscript and software.

