## Supporting Information for "ProteoScopeR: A Shiny Workflow for Method Comparison and Reproducible Quantitative Proteomics Analysis"

Benbo Gao and Hanqing Zhao

#### S1 Software inputs, operations and outputs

Table S1. Programmatic and interactive workflow components.

| Component | Input and operation | Output |
|---|---|---|
| Import | User-selected processor, quantified tables, and metadata; validate and align | ProteoScopeData with assay, annotations, and history |
| Design | Metadata, formula, and contrasts; check rank and estimability | Design and contrast matrices; diagnostic report |
| QC and filtering | Imported assay; inspect missingness, distributions, and detection | QC tables, feature retention, and filtered matrix |
| Normalization | Filtered matrix; compare eligible transformations | Candidate metrics, selected matrix, and plots |
| Missing data | Selected normalized matrix; compare filling and no-filling strategies | Filled and incomplete matrices; sensitivity summaries |
| Differential analysis | Matrix, design, and contrasts; fit selected engines | Common result columns, issue flags, and agreement |
| Optional AI review | Export evidence, import proposals, record decisions, and lock | Recommendation, scientist audit, and execution receipt |
| Export | Completed analysis state; align reusable objects | Native bundle, R objects, and xOmicsShiny objects |

Core operations are callable without Shiny. The interface adds interaction and visualization; it does not define a separate statistical estimator. The Shiny input page does not automatically select a platform: the processor chosen by the user is used, and the submitted files are validated against that route. Generic programmatic imports may still use file-format and column-schema recognition.

#### S2 Exact v1.0.0 case-study configuration

The input contained 3,667 protein groups and 69 samples. Overall detection ≥ 0.30 retained 3,662 groups. All samples were retained. The independent design was ~ 0 + group, with ordered groups pmCNV, nAMD, Ctrl, and PM and contrasts nAMD–Ctrl, pmCNV–PM, and nAMD–pmCNV. No age or sex terms were included. The design had rank four and 65 residual degrees of freedom.

Table S2. Engine routing in the locked ProteoScopeR v1.0.0 analysis. All branches use the same filtering, cyclic-loess normalization, and group-only specification.

| Engine | Matrix | Missing entries | Model details |
|---|---|---|---|
| limma | Canonical MinProb-imputed | 0 | lmFit, contrasts.fit, and empirical Bayes with trend = TRUE |
| Linear model | Same complete matrix as limma | 0 | Ordinary least squares; 65 residual degrees of freedom |
| proDA | Normalized incomplete | 43,836 | Probabilistic dropout model; no preceding imputation |
| msqrob2 | Same incomplete matrix | 43,836 | Robust model; robust = TRUE, ridge = FALSE |
| MSstats | Same incomplete matrix, converted to adapter input | 43,836 before conversion | Protein group as synthetic feature; condition and biological replicate; no added covariates |

For the complete branch, MinProb was generated by imputeLCMD::impute.MinProb with q = 0.01, tune.sigma = 1. The MSstats converts protein-group $\log_2$ abundance to positive intensity and uses synthetic peptide, charge, and fraction fields; it is not a precursor-level MSstats workflow. Matching the starting matrix and group terms does not make the internal estimators identical.

The review package supplies six-method normalization and seven-strategy missing-data summaries. Quick effects are nAMD-minus-pmCNV mean differences and the screen uses Welch tests with BH adjustment; these are not the four-group fitted-engine selections.

Table S3. Seven-strategy missing-data screen after cyclic-loess normalization. Filled-value location is the median imputed value minus the observed 10th percentile.

| Strategy | Retained | Missing % | Filled location | Effect r | Screen selected |
|---|---|---|---|---|---|
| None | 3,662 | 17.35 | N/A | 1.000 | 13 |
| Complete case | 1,025 | 0.00 | N/A | 1.000 | 58 |
| MinProb | 3,662 | 0.00 | -1.515 | 0.679 | 14 |
| KNN | 3,662 | 0.00 | 0.776 | 0.924 | 17 |
| missForest | 3,662 | 0.00 | 0.847 | 0.958 | 15 |
| QRILC | 3,662 | 0.00 | -2.411 | 0.572 | 13 |
| No filling / proDA | 3,662 | 17.35 | N/A | 1.000 | 13 |

The MinProb row is the canonical v1.0.0 calculation. KNN used the impute package, missForest used the missForest package, and QRILC used imputeLCMD::impute.QRILC, and no-filling/proDA is unchanged input in this preprocessing screen rather than a proDA fit. Complete-case agreement uses its retained subset, whose smaller BH testing family changes the comparison.

### S3 Testability, selections, and overlap

Testability requires a finite estimate and adjusted P value. Selection additionally requires adjusted $P < 0.05$ and $|\text{estimate}| \geq 0.5$. The common-testable universe contains 3,662 groups for nAMD–pmCNV, 3,659 for pmCNV–PM, and 3,661 for nAMD–Ctrl. Adjusted P values were not recalculated after intersection.

Table S4. Pairwise nAMD–pmCNV agreement on 3,662 jointly testable groups. Only A/B counts denote exclusive selections in that pair.

| A / B | Effect r | Shared | Only A | Only B | Jaccard |
|---|---|---|---|---|---|
| limma / proDA | 0.805 | 31 | 23 | 6 | 0.517 |
| limma / MSstats | 0.677 | 46 | 8 | 22 | 0.605 |
| limma / msqrob2 | 0.666 | 48 | 6 | 65 | 0.403 |
| limma / Linear model | 1.000 | 52 | 2 | 2 | 0.929 |
| proDA / MSstats | 0.955 | 37 | 0 | 31 | 0.544 |
| proDA / msqrob2 | 0.935 | 37 | 0 | 76 | 0.327 |
| proDA / Linear model | 0.805 | 31 | 6 | 23 | 0.517 |
| MSstats / msqrob2 | 0.979 | 66 | 2 | 47 | 0.574 |
| MSstats / Linear model | 0.677 | 48 | 20 | 6 | 0.649 |
| msqrob2 / Linear model | 0.666 | 50 | 63 | 4 | 0.427 |

Thirty-one groups were selected by all five workflows for nAMD–pmCNV, none for pmCNV–PM, and four for nAMD–Ctrl. These intersections describe agreement, not a ground-truth positive set.

For pmCNV–PM, the MSstats and msqrob2 each had non-estimable effects for A0A5C2GFQ6, A0A5C2GNU8, and Q15286. For nAMD–Ctrl, each had a non-estimable effect for P11597. The MSstats encoded these effects as infinity with missing raw P values and oneConditionMissing flags; the finite-effect rule excludes them. msqrob2 reported missing tests. The diagnostic text remains in differential_results.csv.

### S4 AI evidence provenance and governance

The review input comprises instructions, a response schema and example, the review configuration, privacy notice, experiment and data dictionaries, design and contrast diagnostics, QC and filtering summaries, six-method normalization comparison, seven-strategy missing-data comparison, and outlier evidence. The submitted workflow recommends cyclic loess, no filling with proDA, all five engines, and investigation of four flagged samples.

The scientist accepted cyclic loess, retained MinProb as the selected missing-data method for comparison with all five engines, and retained the four flagged samples after manual checks. The audit records the modified decisions and rationales. The recommendation does not record model provider, model name/version, generation date, or settings; these remain explicitly 'not recorded'.

There is no integrated external model API. Data leave the local application only if a user downloads and transfers a package. Pseudonymization and metadata minimization reduce disclosure but do not constitute anonymization. Review IDs bind recommendations to a run; structural validation checks supported methods, response types, evidence fields, and sample identity, but cannot establish factual correctness. Recommendations are never evaluated as R code, and scientist approval remains required.

### S5 Additional comparison figures

Figure S1. Normalization sensitivity metrics for the six candidates. Mean–variance correlation and quick-screen selections are descriptive diagnostics, not accuracy rankings.

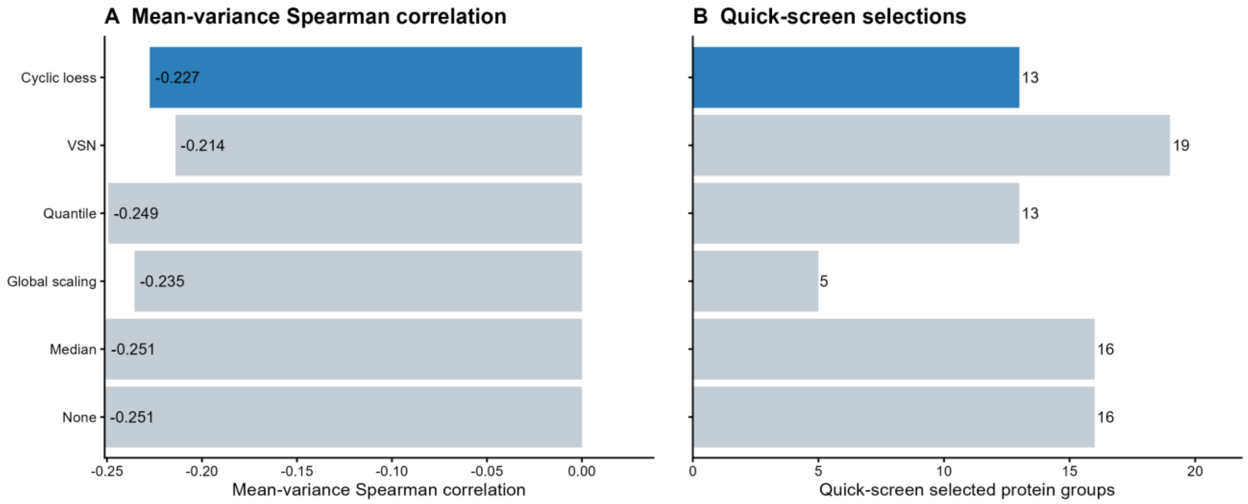


Figure S2. Median within-group coefficient of variation and median sample correlation for the six candidates. The CV is calculated after exponentiation and is scale-dependent, particularly after VSN.

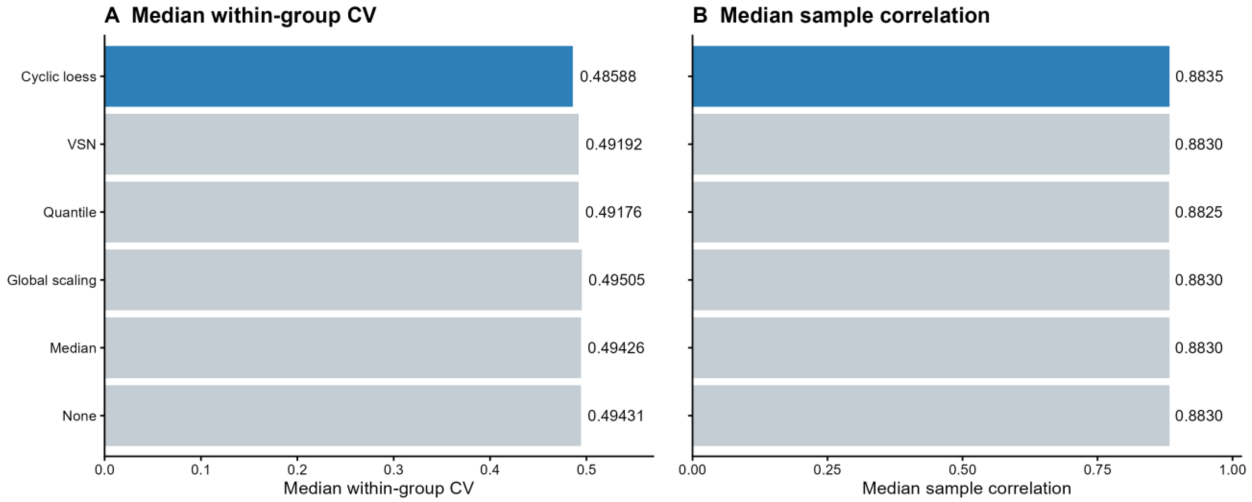


Figure S3. Seven-strategy missing-data sensitivity summary after cyclic loess. The MinProb row is canonical imputeLCMD::impute.MinProb (q = 0.01, tune.sigma = 1).

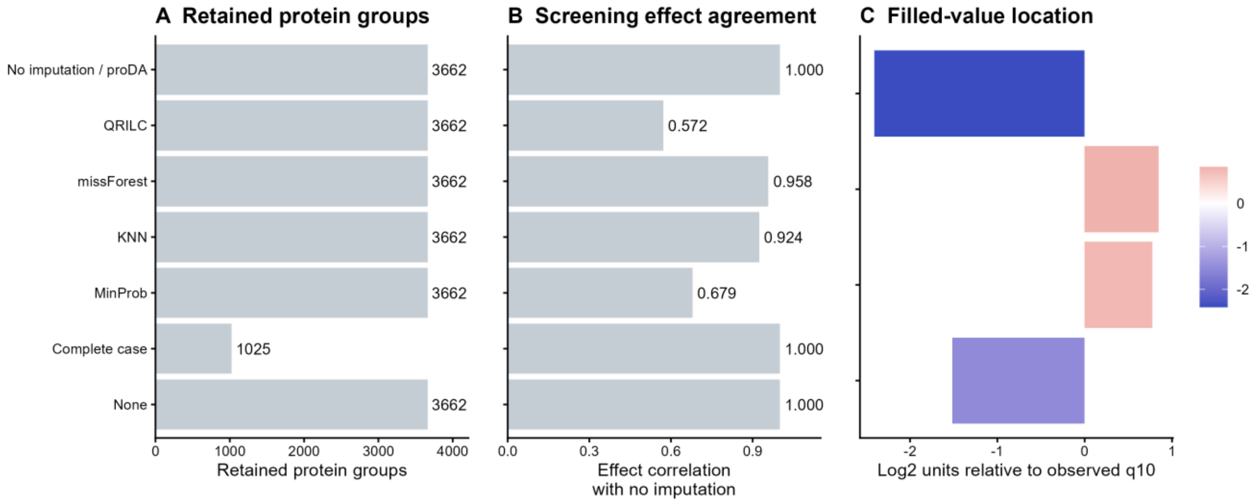


Figure S4. pmCNV–PM volcano comparison using finite results from the locked v1.0.0 analysis. Selected counts are limma 3, proDA 0, MSstats 4, msqrob2 10, and linear model 2. Original engine–contrast adjustments are retained. DEP denotes the number of protein groups meeting both selection thresholds in each panel.

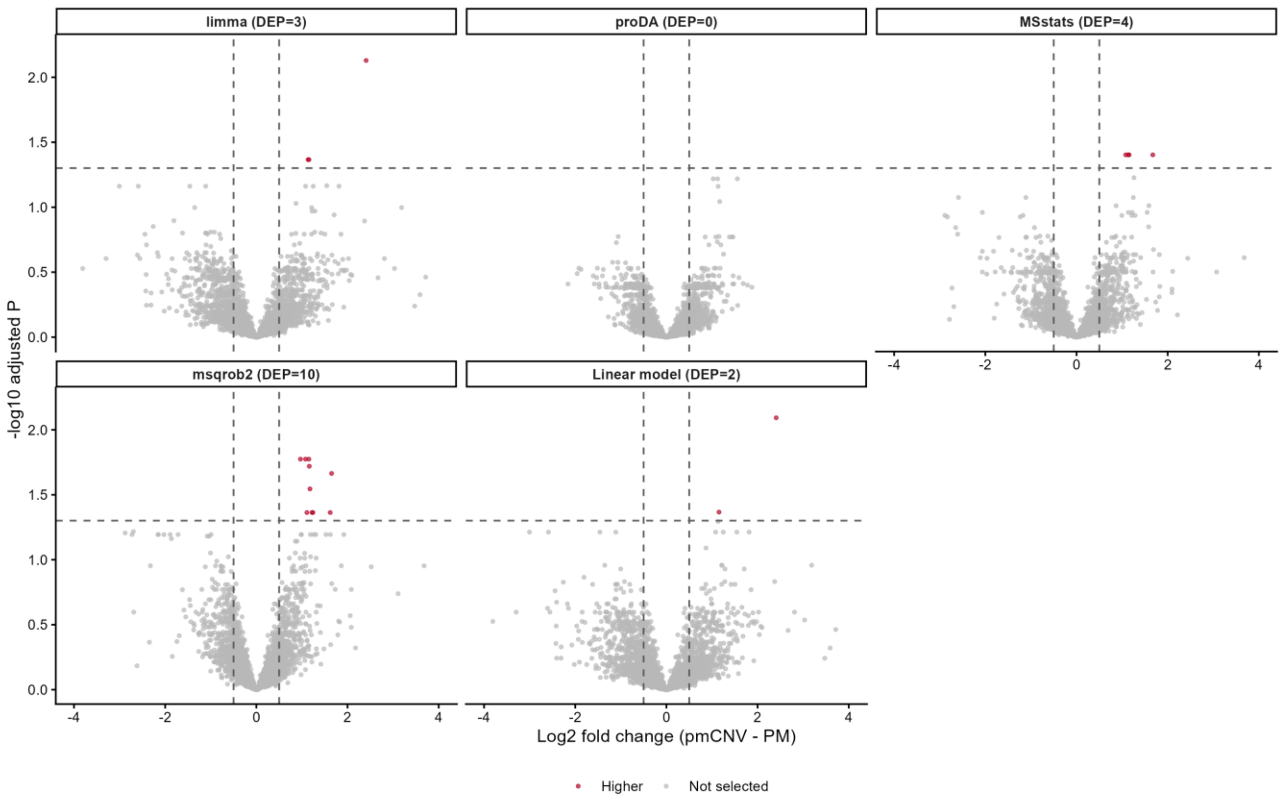
limma (DEP=3)
proDA (DEP=0)
MSstats (DEP=4)
msqrob2 (DEP=10)
Linear model (DEP=2)
-log10 adjusted P
Log2 fold change (pmCNV - PM)
Higher
Not selected

74
